# Local conjugation of auxin by the GH3 amido synthetases is required for normal development of roots and flowers in Arabidopsis

**DOI:** 10.1101/2021.10.04.463114

**Authors:** Ruipan Guo, Yun Hu, Yuki Aoi, Hayao Hira, Chennan Ge, Xinhua Dai, Hiro Kasahara, Yunde Zhao

## Abstract

Gretchen Hagen 3 (GH3) amido synthetases conjugate amino acids to a carboxyl group of small molecules including hormones auxin, jasmonate, and salicylic acid. The Arabidopsis genome harbors 19 *GH3* genes, whose exact roles in plant development have been difficult to define because of genetic redundancy among the *GH3* genes. Here we use CRISPR/Cas9 gene editing technology to delete the Arabidopsis group II *GH3* genes, which are able to conjugate indole-3-acetic acid (IAA) to amino acids. We show that plants lacking the eight group II *GH3* genes (*gh3 octuple* mutants) accumulate free IAA and fail to produce IAA-Asp and IAA-Glu conjugates. Consequently, *gh3 octuple* mutants have extremely short roots, long and dense root hairs, and long hypocotyls and petioles. Our characterization of *gh3 septuple* mutants, which provide sensitized backgrounds, reveals that *GH3*.*17* and *GH3*.*9* play prominent roles in root elongation and seed production, respectively. We show that *GH3* functions correlate with their expression patterns, suggesting that local deactivation of auxin also contributes to maintaining auxin homeostasis and is important for plant development. Moreover, this work provides a method for elucidating functions of individual members of a gene family, whose members have overlapping functions.

## Introduction

Auxin is essential for many aspects of plant development (Zhao, 2018). Auxin concentrations in plants need to be precisely controlled to ensure optimal growth and development in response to environmental and developmental signals. Plants have evolved sophisticated mechanisms to maintain auxin homeostasis. Cellular auxin concentrations are affected by auxin biosynthesis, polar transport, degradation, and conjugation to various molecules including amino acids and sugars (Woodward and Bartel, 2005; Zhao, 2010). Both local auxin biosynthesis and polar auxin transport have been extensively studied and they are required for various plant developmental processes. Disruption of local auxin biosynthesis leads to defects in embryogenesis, seedling growth, vascular development, and flower development (Cheng et al., 2006; Cheng et al., 2007; Stepanova et al., 2008; Tao et al., 2008). Defective polar auxin transport causes characteristic phenotypes such as the formation of pin-like inflorescences (Galweiler et al., 1998). In contrast, much less is understood regarding the roles of auxin degradation and conjugation in plant development.

The *DAO* gene in rice encodes a 2-oxoglutarate and Fe(II) dependent dioxygenase that is able to oxidize indole-3-acetic acid (IAA), the main natural auxin, into 2-oxindole-3-acetic acid (Zhao et al., 2013). Mutations in *DAO* in rice affects several reproductive processes such as anther dehiscence, pollen fertility, and seed initiation (Zhao et al., 2013). It has been shown that DAO plays a similar role in auxin oxidation in Arabidopsis, but the *dao* mutants in Arabidopsis only displayed subtle developmental phenotypes (Mellor et al., 2016; Porco et al., 2016; Zhang et al., 2016). *Gretchen Hagen 3* (*GH3*) genes (Wright et al., 1987), which encode amido synthetases that catalyze the formation of an amide bond between a carboxyl group and the amino group of an amino acid (Staswick et al., 2005), were proposed to compensate for DAO functions in Arabidopsis (Zhang and Peer, 2017). Arabidopsis genome has 19 *GH3* genes, which can be divided into three groups on basis of sequence similarities and gene structure (Okrent and Wildermuth, 2011). Group I is composed of *GH3*.*10* and *GH3*.*11* (Okrent and Wildermuth, 2011). *GH3*.*11* (*JAR1*) catalyzes the conjugation of jasmonate (JA) to isoleucine to form jasmonoyl-isoleucine (JA-Ile) conjugate, which is the active molecule perceived by JA receptors (Staswick and Tiryaki, 2004). The *jar1* mutants were insensitive to methyl jasmonate and were more susceptible to pathogens (Staswick et al., 2002). The functions of *GH3*.*10* are not understood, but overexpression of *GH3*.*10* leads to short hypocotyl under red light conditions (Takase et al., 2003). The group II *GH3* genes in Arabidopsis consists of eight members (Okrent and Wildermuth, 2011). All of the group II GH3 proteins have been experimentally demonstrated to be capable of conjugating auxin to amino acids (Okrent et al., 2009; Westfall et al., 2016). Moreover, the expression of *GH3*.*1, GH3*.*3, GH3*.*5*, and *GH3*.*6* is induced by auxin, suggesting that the group II *GH3* genes have important functions in maintaining auxin homeostasis (Okrent and Wildermuth, 2011). The group III *GH3* has nine members in Arabidopsis and most of the *GH3* in this group have not been functionally or biochemically characterized. The expression of *GH3*.*12* is induced by salicylic acid (SA) and GH3.12 can use benzoate, an SA analog, as a substrate (Okrent and Wildermuth, 2011). Moreover, *gh3*.*12* mutants displayed SA-related phenotypes (Okrent et al., 2009). Recently, GH3.12 (PBS3) was discovered to conjugate isochorismate with glutamate to produce isochorismate-glutamate, which is non-enzymatically and spontaneously converted into SA (Rekhter et al., 2019). GH3.15 uses indole 3-butyric acid and glutamine as substrates in vitro, suggesting that GH3.15 may also participate in auxin homeostasis (Sherp et al., 2018).

Genetic dissection of *GH3* functions has been difficult because of the genetic redundancy among the large number of *GH3* genes in Arabidopsis. Gain-of-function studies of *GH3* genes have clearly suggested that *GH3* genes likely play important roles in plant development. For example, overexpression of *GH3*.*2* in Arabidopsis (*ydk1-D* mutants) leads to short primary roots, few lateral roots, and dwarf plants (Takase et al., 2004). The *dwarf in light 1* (*dfl1-D*) mutants, which overexpress *GH3*.*6*, have short hypocotyls and is extremely dwarf (Nakazawa et al., 2001). Overexpression of *GH3*.*5* in *wes1-D* mutants leads to very short hypocotyls (Park et al., 2007b). Single loss-of-function *gh3* mutants in Arabidopsis only display subtle phenotypes in hypocotyl elongation, primary root development, and lateral root initiation (Khan and Stone, 2007; Porco et al., 2016; Zheng et al., 2016). Another approach for analyzing *GH3* functions was to generate knockout mutants in plants that have fewer *GH3* genes than Arabidopsis. The moss *Physcomitrella patens* only has two *GH3* genes. Moss mutants without the two *GH3* genes have elevated free IAA concentrations and a decreased level of amide-IAA conjugates (Bierfreund et al., 2004; Ludwig-Muller et al., 2009). Moreover, the double mutants were more sensitive to exogenous IAA. However, the double mutants appeared to develop normally under laboratory growth conditions (Bierfreund et al., 2004; Ludwig-Muller et al., 2009).

In order to assess the roles of *GH3* genes in auxin homeostasis and plant development, we used CRISPR/Cas9 gene editing technology to generate true knockout mutants for each of the group II *GH3* genes in Arabidopsis in the Columbia background. The single *gh3* mutants did not display dramatic developmental defects. However, Arabidopsis plants with all group II *GH3* genes inactivated (*gh3 octuple*) display strong auxin over accumulation phenotypes. Light grown *gh3 octuple* mutants have extremely short primary roots, long lateral roots, dense and long root hairs, and much elongated petioles. Dark-grown *gh3 octuple* seedlings have short hypocotyls. We are able to assess the relative contributions of individual *GH3* genes in auxin homeostasis and Arabidopsis development by characterizing several *gh3 septuple* mutants, which we believe provide sensitized backgrounds for analyzing the roles of auxin conjugation. We show that *GH3*.*17* plays a predominant role in controlling root elongation whereas *GH3*.*9* is the main player in controlling Arabidopsis fertility. Moreover, functions of *GH3* genes appear to correlate with the expression patterns of *GH3* genes, suggesting that local auxin conjugation plays key roles in maintaining auxin homeostasis and plant development.

## Results

### Generation of new null mutants for *GH3 amido synthetase* genes

The group II *GH3* family in Arabidopsis consists of 8 genes (*GH3*.*1, GH3*.*2, GH3*.*3, GH3*.*4, GH3*.*5, GH3*.*6, GH3*.*9*, and *GH3*.*17*), which can be divided into three sub-groups (Figure 1A) based on their genomic structures, sequence homology, and the number of intron/exons. GH3 proteins are also more closely related to their sub-group members than to members in a different sub-group (Figure 1B). In order to analyze the roles of *GH3* genes in auxin metabolism and in plant development, we took advantage of the recently developed CRISPR/Cas9 gene editing technology to generate new null alleles for all of the group II *GH3* genes in Arabidopsis (Figure S1). All of the mutants were generated using two guide RNAs, which led to a deletion of a large fragment of a *GH3* gene. Because the coding regions of *GH3*s are largely deleted, the resulting *gh3* mutants are null. Another advantage of using double guide RNAs in editing *GH3* genes is that the mutants can be easily genotyped using PCR-based methods (Table S1).

**Figure 1.**
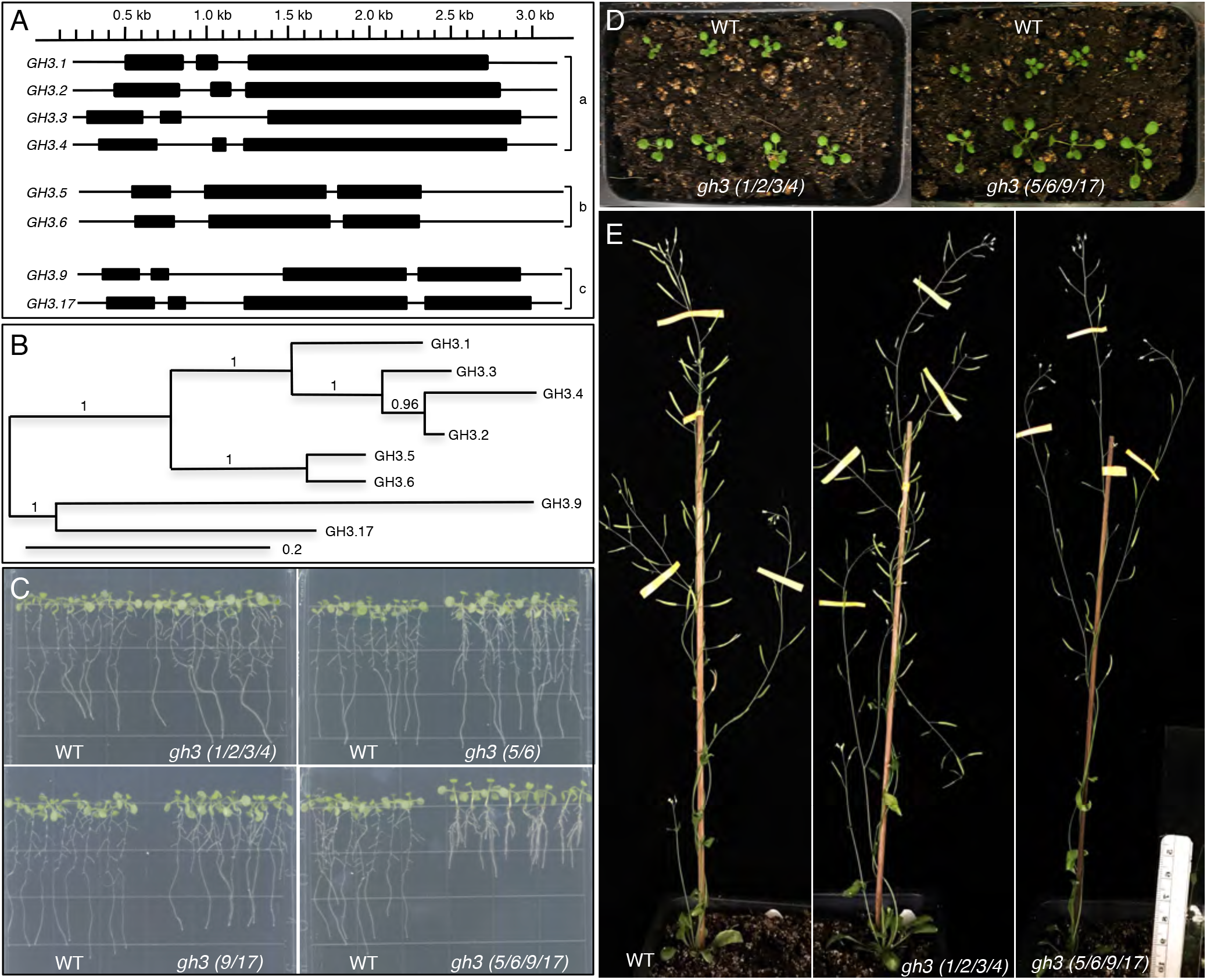
Roles of *GH3* family of genes in Arabidopsis development. A) The Group II *GH3* genes, which encode enzymes capable of synthesizing IAA-amino acid conjugates and/or which are induced by auxin. The *GH3*s can be divided into three sub-groups based on their exon/intron structures. B) A phylogenetic tree of the Arabidopsis Group II GH3 proteins. Note that GH3.1 to GH3.4 form a clade whereas the other GH3s form two other distinct clades. C) Disruption of *GH3* genes affect root development and seedling growth. Simultaneously disruption of *GH3*.*1, GH3*.*2, GH3*.*3*, and *GH3*.*4* (The quadruple mutants are named *gh3 (1/2/3/4)*) leads to slightly longer roots and more lateral roots than WT plants. The *gh3 (5/6/9/17)* had much shorter roots and longer hypocotyls than WT plants. C) The *GH3* genes play a role in petiole elongation. Both *gh3 (1/2/3/4)* and *gh3 (5/6/9/17)* have longer petioles than WT plants. E) *GH3* genes are important for Arabidopsis fertility. Note that *gh3 (5/6/9/17)* have a much reduced fertility.

We analyzed the single *gh3* mutants (at least two independent alleles of each gene) grown under normal laboratory growth conditions. The single mutants did not display dramatic developmental defects. The lack of strong phenotypes of the *gh3* single mutants was likely caused by genetic redundancy among the *GH3* genes. We next constructed multiple mutants on basis of sequence homology of the GH3 proteins. We generated *gh3(1/2/3/4)* quadruple mutants in which *GH3*.*1, GH3*.*2, GH3*.*3*, and *GH3*.*4* had been deleted. We also generated *gh3(5/6)* double mutants, *gh3 (9/17)* double mutants, and *gh3(5/6/9/17)* quadruple mutants (Figure 1C, Figure S2). We generated the double and quadruple mutants by crossing the single mutants together.

Simultaneous disruption of *GH3*.*1, GH3*.*2, GH3*.*3, GH3*.*4* led to slightly longer primary roots than WT (Figure 1C). The *gh3(1/2/3/4)* quadruple mutants also developed more lateral roots (Figure 1C, Figure S2). The *gh3(5/6)* double mutants were very similar to WT except that the double mutants had more lateral roots (Figure 1C, Figure S2). Both *gh3(9/17)* double mutants and *gh3(5/6/9/17)* quadruple mutants had shorter primary roots than WT plants (Figure 1C, Figure S2). The primary roots of *gh3(5/6/9/17)* quadruple mutants were only half the length of those of WT plants (Figure 1C, Figure S2). The *gh3(5/6/9/17)* quadruple mutants also had longer lateral roots than both WT and the *gh3(9/17)* double mutants (Figure 1C). At young adult stages, both *gh3(1/2/3/4)* and *gh3(5/6/9/17)* had longer petioles (Figure 1D, Figure S2), but the *gh3(5/6/9/17)* had a more pronounced phenotype than *gh3 (1/2/3/4)*. Fertility of *gh3(5/6/9/17)* was greatly reduced (Figure 1E). It is clear that *GH3*.*5, GH3*.*6, GH3*.*9*, and *GH3*.*17* play a more prominent role in Arabidopsis development than the other group II *GH3* genes.

### A complete removal of the group II *GH3* genes causes dramatic developmental defects

We generated *gh3 octuple* mutants by crossing the *gh3(5/6/9/17)* quadruple mutants to *gh3(1/2/3/4)* quadruple mutants and genotyped the offspring of the next few generations. The *gh3 octuple* mutants displayed extremely short primary roots (Figure 2A, Figure S3). The *gh3 octuple* mutants also had longer and more densely packed lateral roots (Figure 2A). Moreover, the *gh3 octuple* mutants had more and longer root hairs than WT plants (Figure 2B). Interestingly, the *gh3 octuple* mutants had shorter hypocotyl and less-prominent apical hook when grown in the dark. (Figure 2C & 2D). Dark-grown seedlings of the previously characterized auxin overproduction mutants such as *sur1* and *yuc1-D* also developed shorter hypocotyls and lacked an apical hook (Boerjan et al., 1995; Zhao et al., 2001), suggesting that *gh3 octuple* mutants might accumulate more auxin than WT. Many of the flowers in *gh3 octuple* mutants aborted before they became mature (Fig 2E). The *gh3 octuple* mutants were almost sterile: *gh3 octuple* mutants hardly produced any pollen grains (Figure 2F). The *gh3 octuple* mutants had much shorter filaments and larger anthers than WT (Figure 2F). The number of floral organs of the *gh3 octuple* mutants was similar to that of WT flowers.

**Figure 2.**
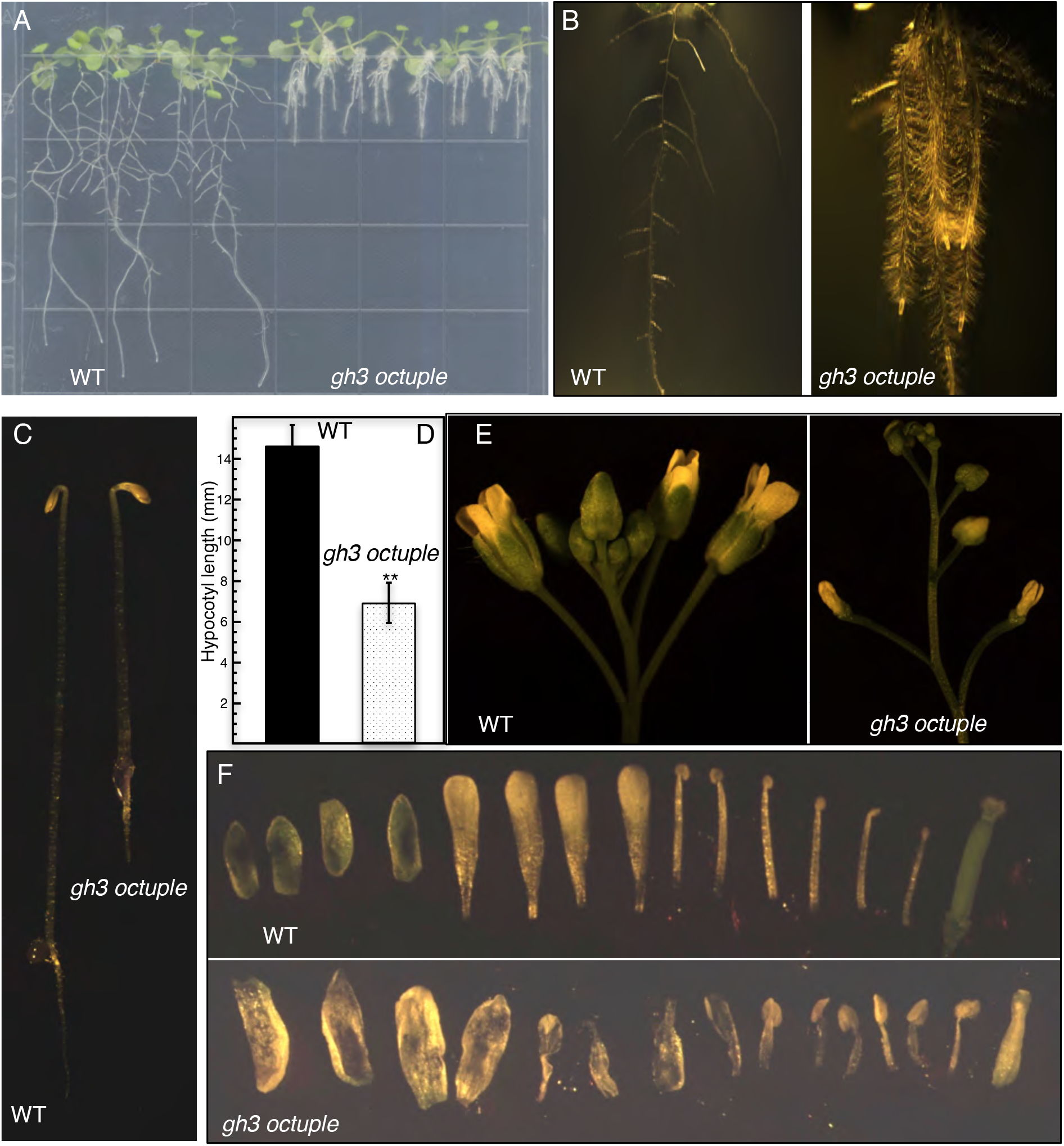
Elimination of group II *GH3* genes impacts major developmental processes in Arabidopsis. A) Arabidopsis plants without the eight group II *GH3* genes (*gh3 octuple*) develop extremely short primary roots and elongated lateral roots. B) The *gh3 octuple* mutants have more and longer root hairs. C &D) Dark-grown *gh3 octuple* plants displayed short hypocotyls. E) Flowers in *gh3 octuple* fail to develop normally. Most *gh3 octuple* flowers abort early without producing any seeds. Left WT, and right *gh3 octuple*. F) Floral defects of *gh3 octuple*. The number of floral organs in *gh3 octuple* mutants is similar to that of WT, but the sexual organs do not develop properly. Note that *gh3 octuple* stamens have enlarged anthers.

### Altered auxin homeostasis in *gh3* mutants

Because the *GH3* proteins conjugate free IAA to amino acids and the IAA-amino acid conjugates are considered inactive, we hypothesized that *gh3 octuple* mutants would accumulate more active auxin. Moreover, the observed developmental defects of *gh3 octuple* mutants were consistent with the hypothesis that *gh3 octuple* mutants accumulated more auxin (Figure 2). It was known that auxin stimulates rooting in tissue culture and that explants of auxin overproduction mutants are able to develop extensive roots in auxin-free media (Zhao et al., 2001). Explants of *gh3 octuple* mutants developed roots in auxin free-media within seven days (Figure 3A, Figure S4). The roots from *gh3 octuple* explants had long and dense root hairs (Figure 3A) whereas WT explants failed to develop roots under the same growth conditions, suggesting that *gh3 octuple* mutants indeed accumulated more auxin. Furthermore, *gh3 octuple* mutants initiated adventitious roots out of hypocotyls (Figure 3B), a characteristic phenotype of known auxin overproduction mutants such as *sur1* and *sur2* (Barlier et al., 2000; Boerjan et al., 1995).

**Figure 3.**
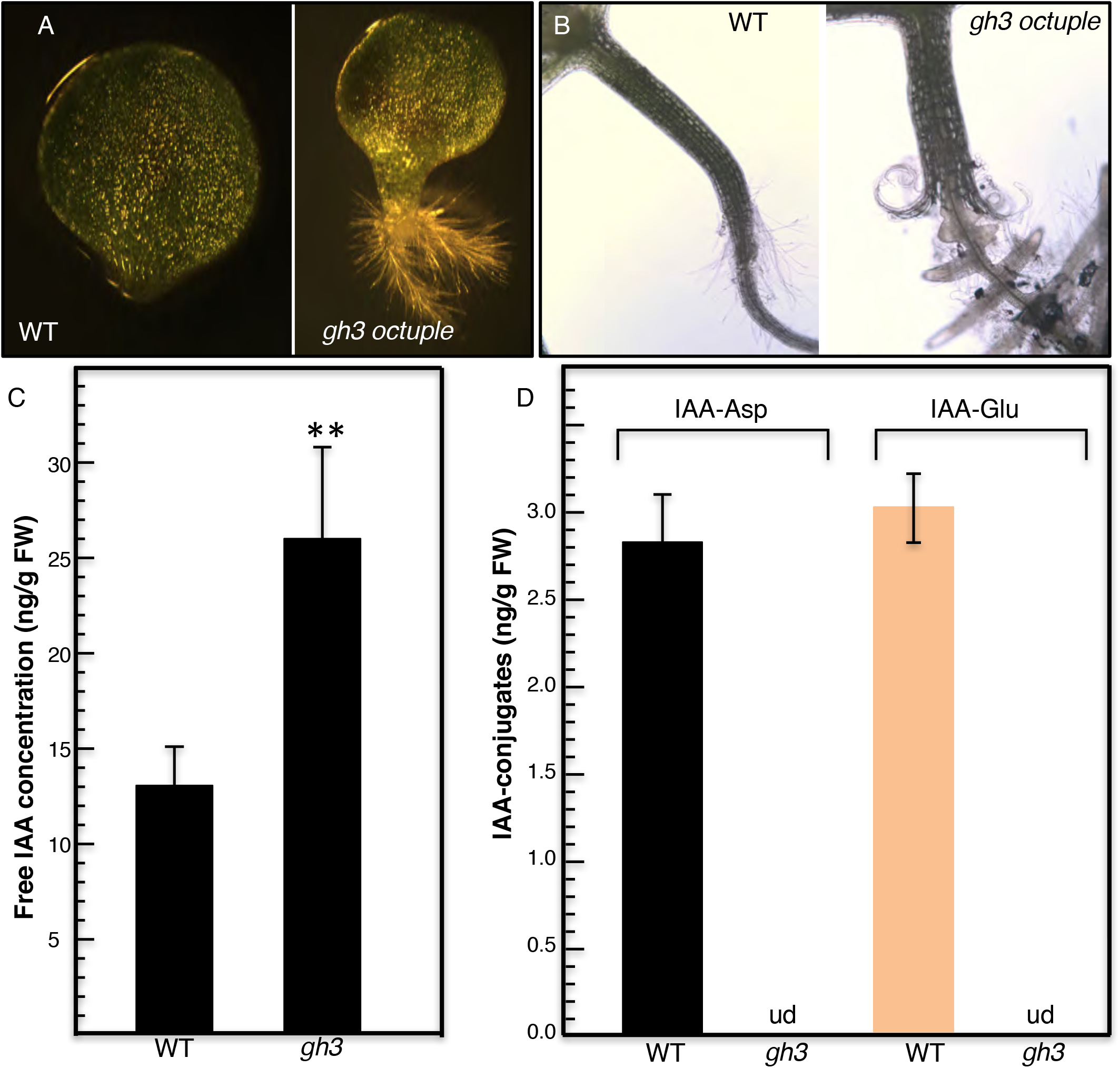
Disruption of *GH3* genes alters auxin homeostasis. A) An explant of *gh3* octuple mutants produced adventitious roots. Explants were grown on hormone free media under light for 7 days. B) Adventitious roots grown out of hypocotyls of *gh3 octuple* mutants. 14-day old seedlings were grown on MS media. C) The *gh3* mutants accumulated more free IAAs than WT. 9-day old seedlings of WT and offspring of *gh3*.*9*^*+/-*^ *Δ7gh3* were grown on vertical MS plates. D) The *gh3* mutants contained un-detectable amount of IAA-Asp and IAA-Glu conjugates. ud: undetectable.

We analyzed the concentrations of free IAA and IAA-Asp and IAA-Glu, two auxin conjugates that are believed inactive, in the *gh3* mutants (Figure 3C & D). Because *gh3 octuple* mutants set very few seeds, we used a segregating population from plants that had a single copy of *GH3*.*9* [*gh3*.*9*^*+/-*^ *gh3 (1/2/3/4/5/6/17)*], which was fertile. As shown in Figure 3C, the *gh3* mutants contained almost 2 times more IAA than WT plants. In addition, both IAA-Asp and IAA-Glu in the *gh3* mutants were below the detection limit of our assay (Figure 3D), suggesting that the *GH3* genes analyzed here are the main contributor to synthesizing IAA conjugates.

### Determination of relative contributions of *GH3* genes to Arabidopsis development

The genetic redundancy among the *GH3* genes makes it difficult to dissect the unique roles of individual *GH3* in auxin homeostasis and plant development. We hypothesized that the *gh3 septuple* mutants might provide sensitized backgrounds for analyzing the functions of individual *GH3* genes. Analyzing plants with only one *GH3* may reveal the functions of the *GH3* in the absence of the interferences from other *GH3* genes. We focus on four *gh3 septuple* mutants that still had one of the *GH3* genes from the subgroups b and c (Figure 1A) because *GH3*.*5, GH3*.*6, GH3*.*9*, and *GH3*.*17* appeared to play a more prominent role based on our analysis of *gh3(1/2/3/4)* and *gh3(5/6/9/17)* quadruple mutants (Figure 1). We name a *septuple*-mutant *Δ7gh3*. The *GH3*.*5Δ7gh3* refers to plants that lacked all group II *GH3* genes except the *GH3*.*5*.

Hypocotyl elongation is very sensitive to changes in auxin concentrations (Collett et al., 2000). The *gh3 octuple* mutants had much longer hypocotyls than WT plants (Figure 4A). Interestingly, both *GH3*.*5 Δ7gh3* and *GH3*.*6 Δ7gh3* had even longer hypocotyls than *gh3 octuple* mutants (Fig 4A, Figure S5). Among the four *gh3 septuple* mutants we analyzed, *GH3*.*5Δ7gh3, GH3*.*6Δ7gh3*, and *GH3*.*17Δ7gh3* had longer roots than the *gh3 octuple* mutants (Figure 4A & 4B, Figure S5), suggesting that *GH3*.*5, GH3*.*6*, and *GH3*.*17* all contribute to root elongation. In contrast, the *GH3*.*9Δ7gh3* and *gh3 octuple* mutants were indistinguishable in terms of root length (Figure 4A &4B, Figure S5). These results suggest that *GH3*.*5, GH3*.*6*, and *GH3*.*17* play a more important role in root development than *GH3*.*9*. Surprisingly, *GH3*.*17* alone was sufficient to reverse the root defects observed in *gh3 octuple* mutants (Figure 4A & 4B). We noticed that *GH3*.*17Δ7gh3* actually had longer primary roots and more lateral roots than WT plants (Figure 4B, Figure S5).

**Figure 4.**
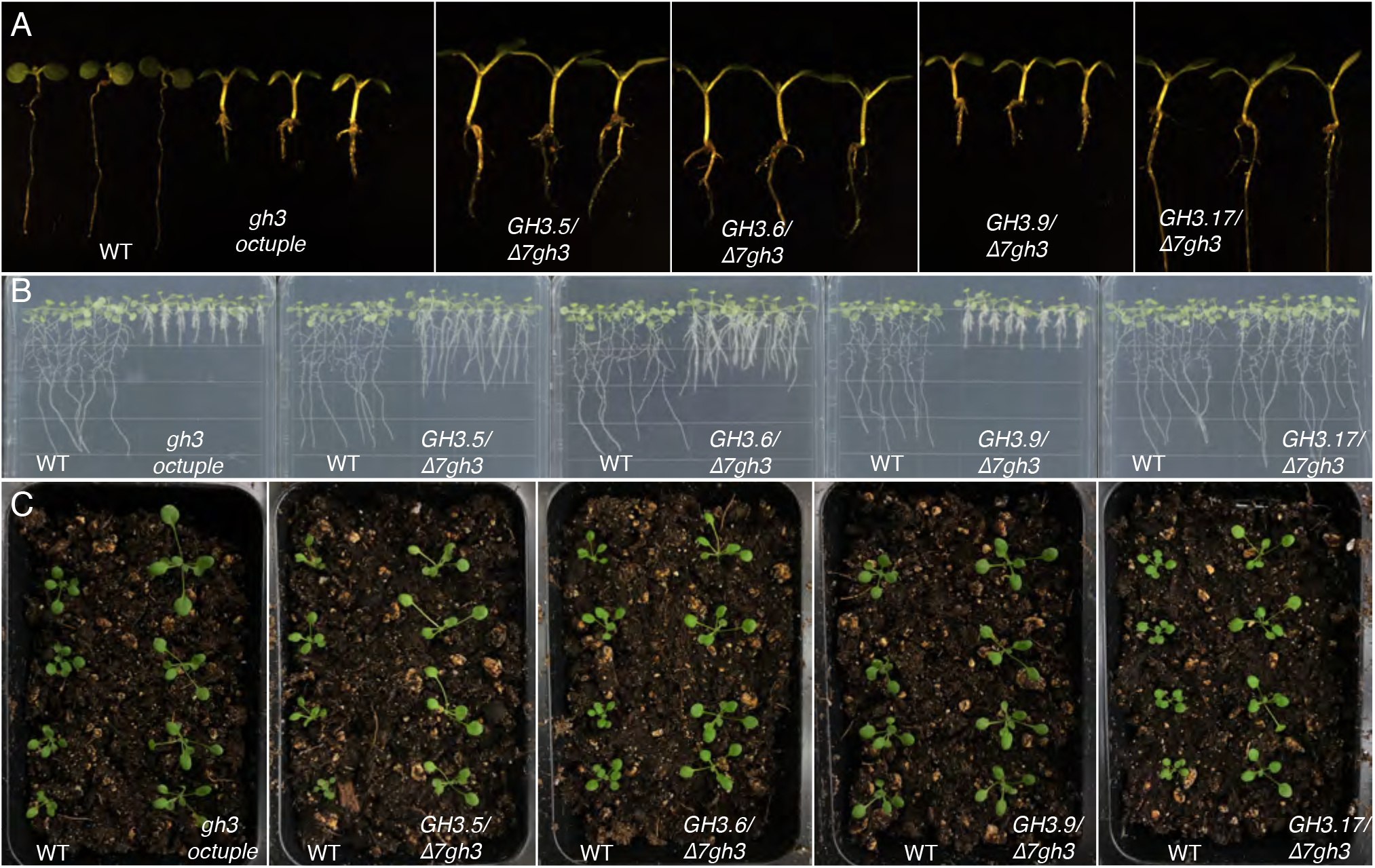
Identification of the prominent GH3 genes responsible for root and seedling development. A) Five-day old light grown seedlings of wt, *gh3 octuple*, and various *gh3 septuple* mutants. The *gh3*.*1 gh3*.*2 gh3*.*3 gh3*.*4 gh3*.*6 gh3*.*9 gh3*.*17 septuple* mutants are named as *GH3*.*5/Δ7gh3*. Note that the septuple mutants differ significantly in terms of hypocotyl length, root length, and cotyledon size. B) *GH3*.*17* plays a more prominent role in root development whereas *GH3*.*9* hardly affects root development. D) Impact of *GH3* genes on petiole elongation. *GH3*.*6/Δ7gh3* refers to plants that lacked all *GH3* genes except GH3.6. *Δ7gh3* refers to plants in which seven of *GH3* genes have been deleted.

The *GH3* genes are important in controlling petiole elongation (Figure 1). A complete deletion of the group II *GH3s* led to much elongated petioles (Figure 4C). Petioles in *gh3 octuple* mutants were more than 2-fold longer than WT (Figure 4C, Figure S3). Compared to *gh3 octuple* mutants, the petiole length in *GH3*.*5Δ7gh3* and *GH3*.*6Δ7gh3* did not change much. In contrast, petioles in *GH3*.*9Δ7gh3* and *GH3*.*17Δ7gh3* were significantly shorter than those in *gh3 octuple* mutants (Fig 4C, Figure S5), suggesting that the two *GH3* genes play a more important role in petiole development than *GH3*.*5* and *GH3*.*6*.

As shown in Figure 2, Arabidopsis plants without the group II *GH3* genes developed abnormal flowers and hardly set any seeds. We compared the fertilities of *gh3 octuple* mutants with those of *gh3 septuple* mutants (Figure 5). All of the *gh3 septuple* mutants except *GH3*.*9Δ7gh3* had very few elongated siliques (Figure 5A). Both *GH3*.*5Δ7gh3* and *GH3*.*6Δ7gh3* had slightly better fertilities compared to *gh3 octuple* (Figure 5A). The floral defects in *GH3*.*5Δ7gh3* and *GH3*.*6Δ7gh3* were not as severe as those in *gh3 octuple* mutants (Fig 5B). Surprisingly, *GH3*.*9Δ7gh3* was as fertile as WT plants with fully elongated siliques (Figure 5A & C).

**Figure 5.**
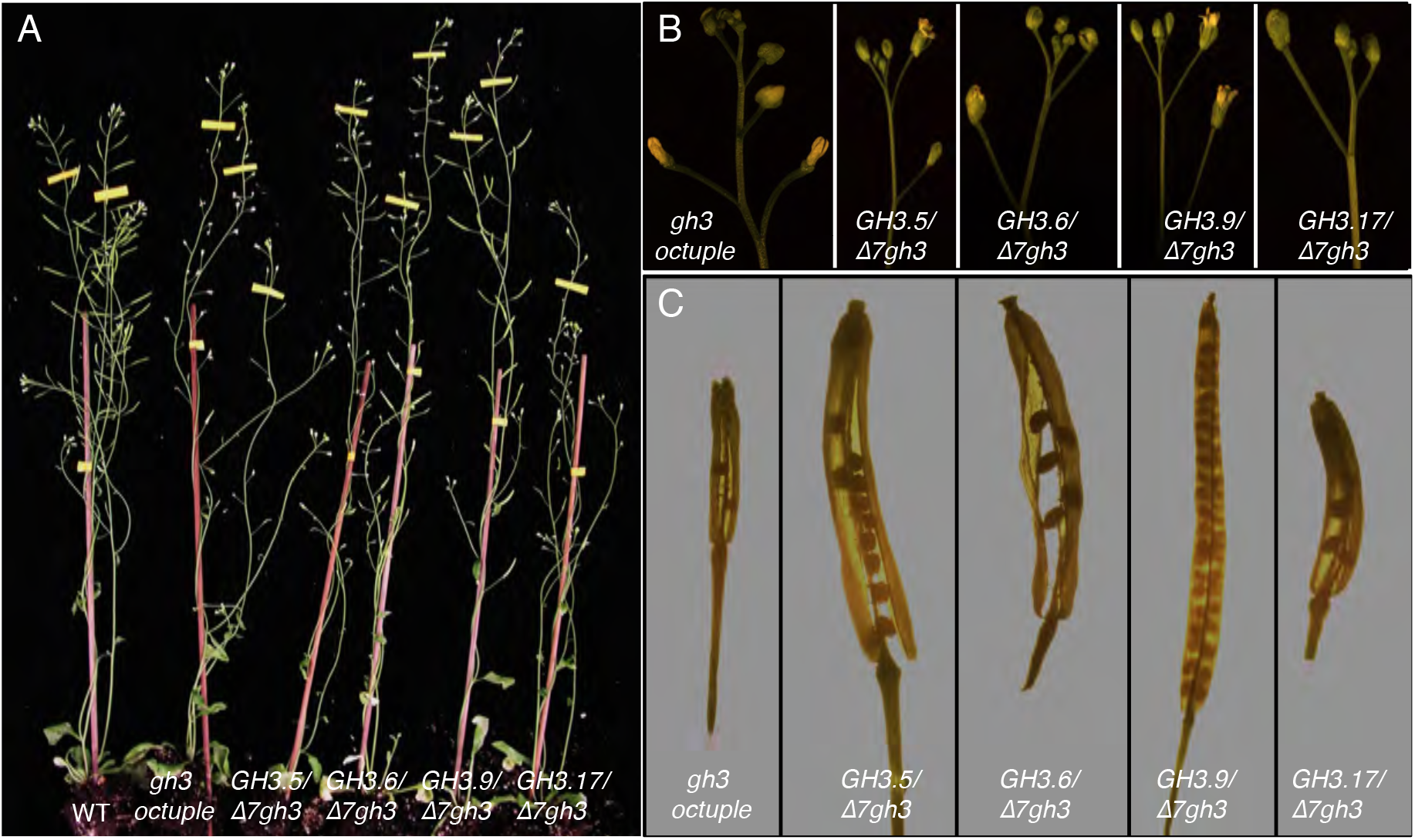
*GH3*.*9* plays a more prominent role in controlling Arabidopsis flower development and fertility. A) *GH3*.*9* alone in the absence of all other group II *GH3* genes (*GH3*.*9/Δ7gh3)* is able to restore Arabidopsis fertility that is abolished by deletion of all *GH3* genes. Note that *gh3 octuple* is almost sterile and *GH3*.*5/Δ7gh3* is able to develop some extended siliques. However, *GH3*.*17/Δ7gh3* and *gh3 octuple* are similarly sterile. B) Inflorescence apex and flowers of *gh3 octuple* mutants and various *gh3 septuple* mutants. C) A comparison of silique length and seed numbers between *gh3 octuple* and *gh3 septuple* mutants

### The *gh3* mutant phenotypes largely correlate with the expression patterns the *GH3*s

We analyzed the expression patterns of *GH3*.*5, GH3*.*6, GH3*.*9*, and *GH3*.*17* using the publically available gene expression profile data (Arabidopsis.org) (Figure S6). The *GH3* genes were hardly expressed in dry seeds, but soaking the seeds in media for one day induced the expression of *GH3*.*5* and *GH3*.*6*, but not the other two *GH3* genes (Figure S6). By the third day of socking, *GH3*.*5, 3*.*6*, and *3*.*17* were highly expressed. Among the four *GH3* genes analyzed, *GH3*.*9* is the only *GH3* that is not highly expressed in roots during seedling stages. The *GH3*.*9* expression patterns were consistent with our observations that *GH3*.*9Δ7gh3* and *gh3 octuple* mutants had similar root defects (Figure 4B). The *GH3* genes that were expressed in roots were able to partially restore root elongation defects of *gh3 octuple* mutants (Figure 4A & 4B). Among the four *GH3* genes, *GH3*.*6* was highly expressed in hypocotyls (Figure S6). Interestingly, *GH3*.*6* had a more pronounced impact on hypocotyl elongation (Figure 4A). *GH3*.*5, GH3*.*6*, and *GH3*.*9* were expressed in flowers and they were able to partially overcome the fertility defects observed in *gh3 octuple* mutants (Figure 5B). In siliques, the highly expressed *GH3* gene was *GH3*.*9* (Figure S6). Consistent with the expression pattern, *GH3*.*9* alone was sufficient to maintain Arabidopsis fertility in the absence of all other group II *GH3* genes (Figure 5)

## Discussion

The roles of GH3 amido synthetases in auxin homeostasis and plant development have been difficult to define because of the genetic redundancy among the *GH3* genes. By generating new null alleles of group II *GH3* genes and by constructing plants without any group II *GH3* genes, we are able to demonstrate that *GH3* genes are required for normal development of Arabidopsis. Plants without the group II *GH3* accumulate more free-IAA and fail to make any IAA-Asp and IAA-Glu, two presumed inactive auxin conjugates. Consequently, *gh3 octuple* mutants have extremely short roots, more and dense root hairs, and long petioles. Moreover, we identify the prominent *GH3* genes responsible for root development and flower development by analyzing *gh3 septuple* mutants. This work not only defines the physiological roles of the group II *GH3* genes, but also provides a method for elucidating the unique functions of members of a gene family, which are often masked by other family members due to genetic redundancy.

Auxin homeostasis is severely disrupted when the group II *GH3* genes are deleted from Arabidopsis plants. The *gh3 octuple* mutants displayed phenotypes that have been observed in plants treated with exogenous auxin and/or auxin overproduction mutants (Figure 2). All auxin overproduction mutants and transgenic lines including *yuc1-D* (Zhao et al., 2001), *iaaM* overexpression lines (Romano et al., 1995), *CYP79B2* overexpression lines (Zhao et al., 2002), *sur1* (Boerjan et al., 1995), and *sur2* (Barlier et al., 2000) have longer hypocotyls than WT plants when grown in light. In darkness, *sur1* and *yuc1-D* have shorter hypocotyls and lacked apical hook (Boerjan et al., 1995; Zhao et al., 2001). Light-grown *gh3 octuple* mutants had much elongated hypocotyls, indicative of high auxin phenotypes (Figure 4). Too much auxin caused by either auxin treatments or auxin overproduction inhibits primary root elongation and stimulates root hair and lateral root development. The *gh3 octuple* mutants had very short primary roots and dense and long root hairs, a phenotype that is likely caused by auxin over accumulation (Figure 2). Our previous studies on *yuc1-D* demonstrated that auxin over accumulation greatly decreases Arabidopsis fertility (Zhao et al., 2001). We observed that the *gh3 octuple* mutants displayed defects in fertility similar to that of *yuc1-D* (Figure 5). All of the developmental defects observed in *gh3 octuple* are consistent with the hypothesis that *gh3 octuple* mutants accumulate more auxin than WT plants. Our analysis of free IAA and IAA-conjugates further confirmed that *gh3* mutants fail to conjugate IAA to Asp and Glu. Consequently, the free IAA concentrations in *gh3* mutants were almost doubled. Our results are consistent with findings from a previous characterization of the sextuple mutants *gh3 (1,2,3,4,5,6)*, which had elevated free IAA, no detectable IAA-Asp, and elevated IAA-Glu (Porco et al., 2016). The difference in IAA-Glu concentrations between our gh3 mutants and the *gh3 (1,2,3,4,5,6)* suggests that GH3.9 and Gh3.17 may prefer to using Glu as a substrate, or that *GH3*.*9* and *GH3*.*17* are more activated in the sextuple mutants or both. Nevertheless, our analysis clearly demonstrated that *GH3*s play a key role in auxin homeostasis and Arabidopsis development. Some of the group II GH3 have previously shown to use SA and JA as substrates (Park et al., 2007a; Westfall et al., 2016; Zhang et al., 2007). Whether some of the phenotypes such as sterility observed in *gh3 octuple* is partially caused by alteration of the homeostasis of other hormones remain to be investigated. A promoter-swap experiment would help determine whether the different developmental defects are caused by the different expression patterns of *GH3* genes or by differences in biochemical properties or both. The genetic materials generated in this work should be very useful for studying hormonal cross-talks.

It is well documented that plant development is very sensitive to changes in auxin concentrations (Collett et al., 2000). Auxin can inhibit or stimulate plant growth depending on the concentrations. Hypocotyl elongation and primary root elongation are two main phenotypic readouts of auxin concentration changes. It is known that auxin overproduction stimulates hypocotyl elongation (Zhao et al., 2001). The *gh3 octuple* mutants had longer hypocotyls than WT (Figure 4). Interestingly, both *GH3*.*5Δ7gh3* and *GH3*.*6Δ7gh3 septuple* mutants had longer hypocotyls than the *gh3 octuple* mutants (Figure 4A). Although we did not analyze the actual IAA concentrations in *GH3*.*5Δ7gh3* and *GH3*.*6Δ7gh3 septuple* mutants, we believe that the two *septuple* mutants likely have accumulated less auxin than *gh3 octuple* mutants. Our results suggest that IAA concentrations in *gh3 octuple* may actually be too high for optimal hypocotyl growth. We also observed that the primary root length of *gh3(1/2/3/4)* and *GH3*.*17Δ7gh3 septuple* mutants was longer than WT (Fig1 &4), suggesting that a slight increase of auxin in Arabidopsis stimulates primary root elongation, though further increases of auxin concentrations inhibit root elongation as shown in *gh3 octuple* mutants. Our results are consistent with previous findings that auxin-dependent plant growth displays a bell-shaped curve.

It is relatively straightforward to demonstrate the overlapping functions among members of a gene family by determining whether the higher order mutants of gene family members have enhanced phenotypes than the lower order mutant combinations. In contrast, determining the unique functions of individual members of a gene family is more complicated because the interference of other family members. We hypothesized that a removal of all gene family members except one may provide sensitized backgrounds for determining functions of the particular member. In contrast to analyzing multiple gene knockouts for phenotypic enhancements, here we determine whether a *GH3* gene is sufficient to suppress a developmental defect displayed in the *gh3 octuple* mutants. We showed that *GH3*.*17* alone was sufficient for restoring root elongation defects observed in the *gh3 octuple* mutants (Figure 4A&B). But *GH3*.*17* had no impact on restoring the fertility of *gh3 octuple* mutants (Figure 5). Interestingly, *GH3*.*9* did not affect root elongation as *GH3*.*9Δ7gh3 septuple* mutants and *gh3 octuple* were almost identical in terms of root elongation (Figure 4B). However, *GH3*.*9* alone was sufficient to reverse the sterile phenotypes observed in *gh3 octuple* mutants. The unique functions of *GH3*.*17* in root elongation and *GH3*.*9* in fertility only became evident in the *septuple* mutants (Figure4 & 5), whereas both *gh3*.*17* and *gh3*.*9* single mutants only had subtle phenotypes. Interestingly, the phenotypes of *GH3*.*17Δ7gh3* and *GH3*.*9Δ7gh3 septuple* mutants correlate with the expression patterns of the two *GH3* genes (Figure S6). It is well known that both local auxin biosynthesis and polar auxin transport are required for generating and maintaining auxin gradients. Local auxin conjugation by the GH3.17/VAS2 was previously shown to play an important role in shade avoidance (Zheng et al., 2016). Our results suggest that local auxin conjugation by GH3.17 is important for root elongation and that auxin conjugation by GH3.9 is critical for Arabidopsis fertility. This work also demonstrated the effectiveness of our using sensitized backgrounds for defining unique functions of individual genes of a gene family, whose members have overlapping functions.

## Materials and method

### Plant materials and Growth Conditions

Arabidopsis thaliana plants used in this study were in the Columbia-0 genetic background. The *gh3* mutants were generated using CRISPR/Cas9 gene editing technology, which was described previously in detail (Gao et al., 2016; Gao et al., 2015; Gao and Zhao, 2014). We used two guide RNA molecules for each *GH3* gene to generate deletions of large fragments. The gRNA target sequences and the locations of the gRNAs are shown in Figure S1. The sizes and the locations of the deletion in each *gh3* mutants are shown in Figure S1 as well. The sequences flanking the deletions are shown in the Figure S1. For example, *gh3-c1* had a 1940 bp deletion and *gh3*.*1-c2* harbored a 1943 bp deletion (Figure S1).

We used a PCR based method to genotype the *gh3* mutants (Gao et al., 2016). A pair of primers that are located outside of the two gRNA target sequences can amplify out a large fragment from WT genomic DNA and a much smaller fragment from a deletion mutant (Figure S1). For example, the GH3.1-GT1 and GH3.1-GT2 primer pair generates a 2.8 kb fragment and a 0.8 kb fragment from WT and *gh3*.*1* mutant DNA, respectively. To further clarify the zygosity of the *gh3* mutants, we designed a third primer that was located between the two gRNA target sequences (Figure S1). The GT3 primer paired with GT1 was used to amplify a fragment from WT DNA, but not from the mutant DNA. The locations and directions of all of the genotyping primers were shown in Figure S1. The primer sequences were listed in Table S1. To generate higher order of *gh3* mutants, we used the *gh3-c1* alleles of each *GH3* gene.

### Measurements of auxin and auxin conjugates

Quantification of IAA and its amino acid conjugates were performed using LC-MS/MS as previously reported (Mashiguchi et al., 2019).

## Acknowledgement

We would like to thank Professor Ken-ichiro Hayashi and members of the Zhao lab for comments and suggestions. This work was supported by NIH grant GM114660 to YZ.

