## Supplemental figures and tables for "Local conjugation of auxin by the GH3 amido synthetases is required for normal development of roots and flowers in Arabidopsis"

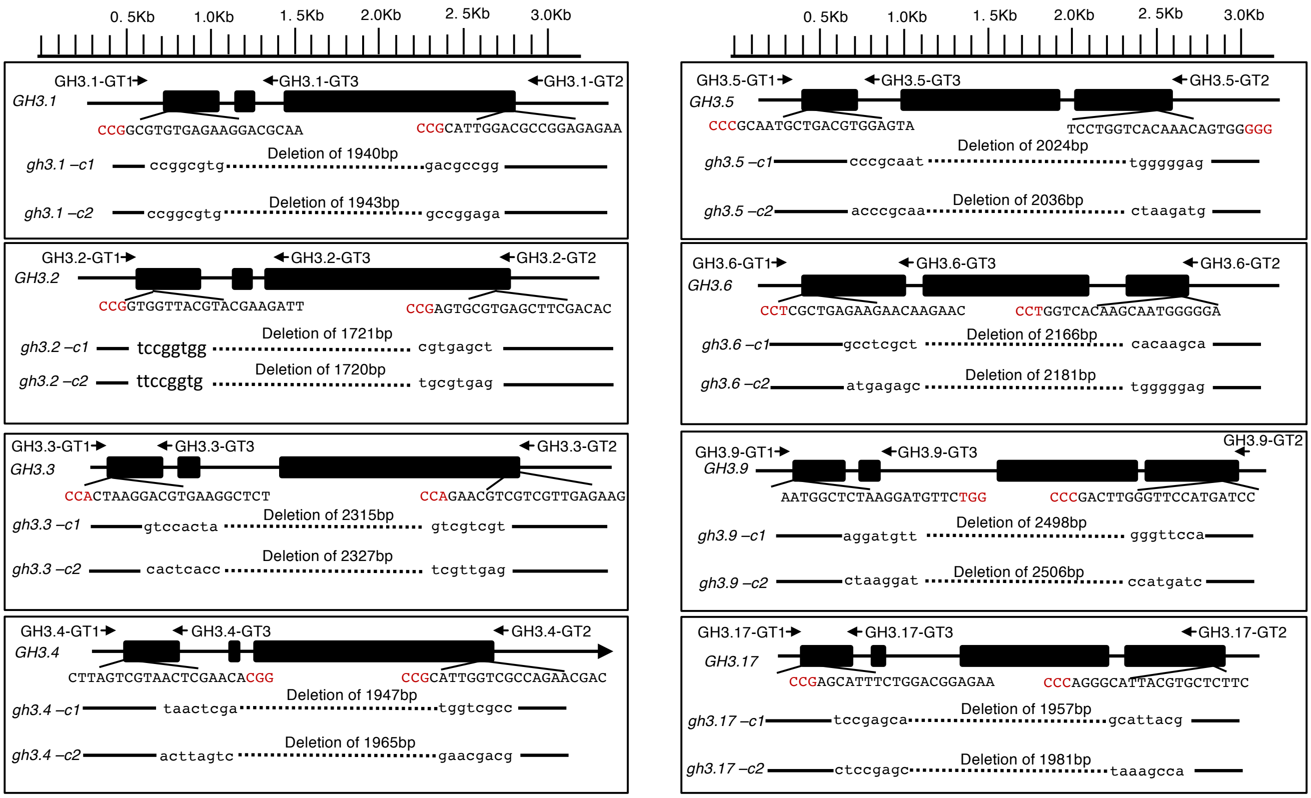


**SI Figure 1. Generation of null mutants for group II *GH3* genes using CRISPR/Cas9 gene editing technology**. The gene structures of each *GH3* gene are schematically shown. Exons are shown in black boxes and introns are indicated by a line. Arrows indicate the directions of genotyping primers. The guide RNA target sequences are shown in upper case with PAM sequences highlighted in red. The sequences flanking the deletions are shown in lower case. The exact sizes of the deletions are shown in the figure. Mutants are identified by PCR using GT1/GT2 primer pairs. Homozygous and heterozygous mutants are differentiated using GT1/GT3 primer pairs.

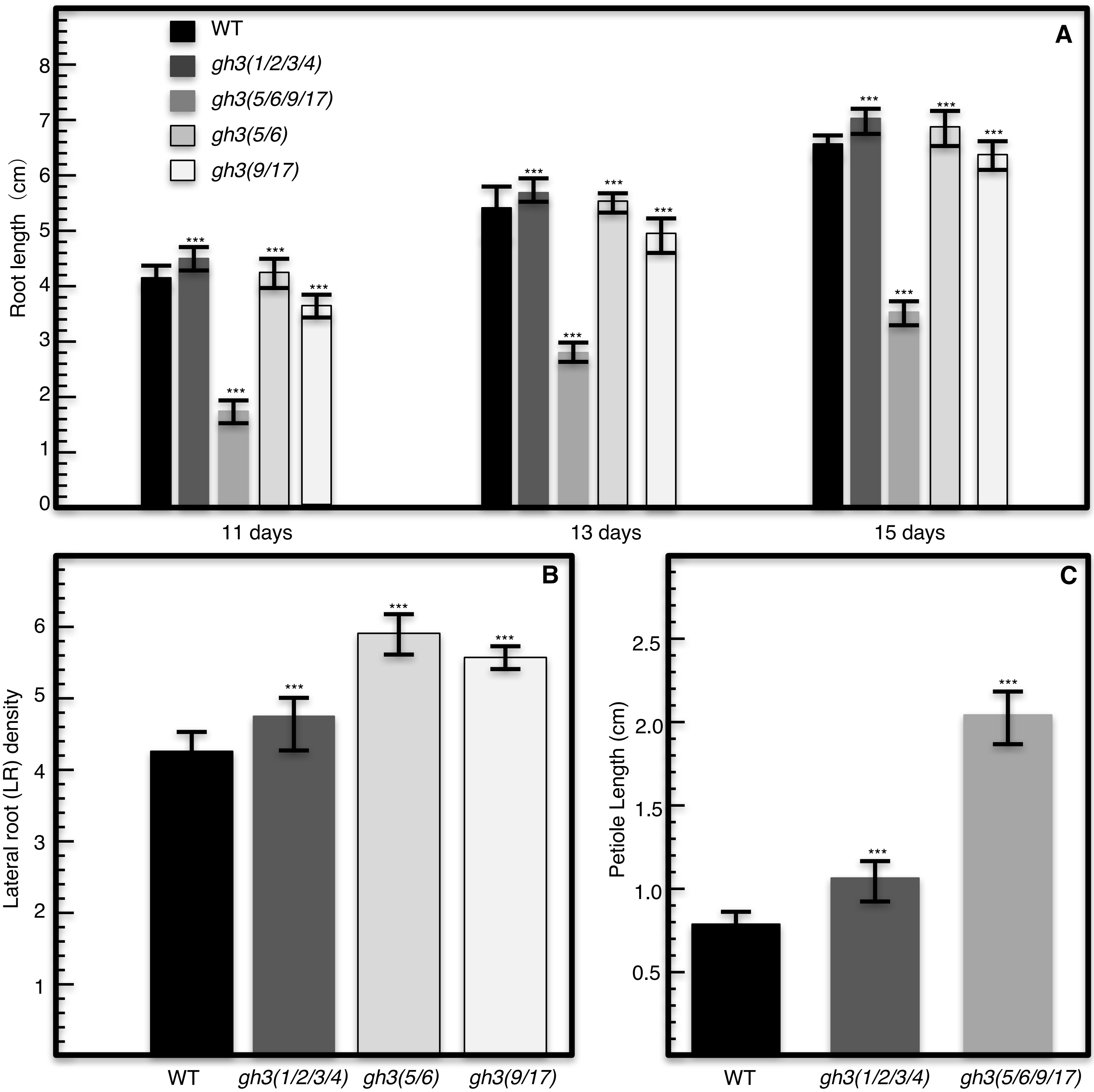


**SI Figure 2. The roles of *GH3* genes in root development and petiole elongation.** A) Primary root length of WT and various *gh3* mutant combinations grown on vertical plates in light for 11, 13, and 15 days. B) Lateral root density in WT and *gh3* mutants. The density was measured as the number of lateral roots per cm primary roots. Samples were 15-day old seedlings grown on vertical plates were analyzed. C) Petiole length of the first true leaf of WT and *gh3* quadruple mutants grown in the greenhouse for two weeks. Twenty seedlings of each genotype were measured for each trait. The results were subjected to student t-test. *** indicates < 0.001.

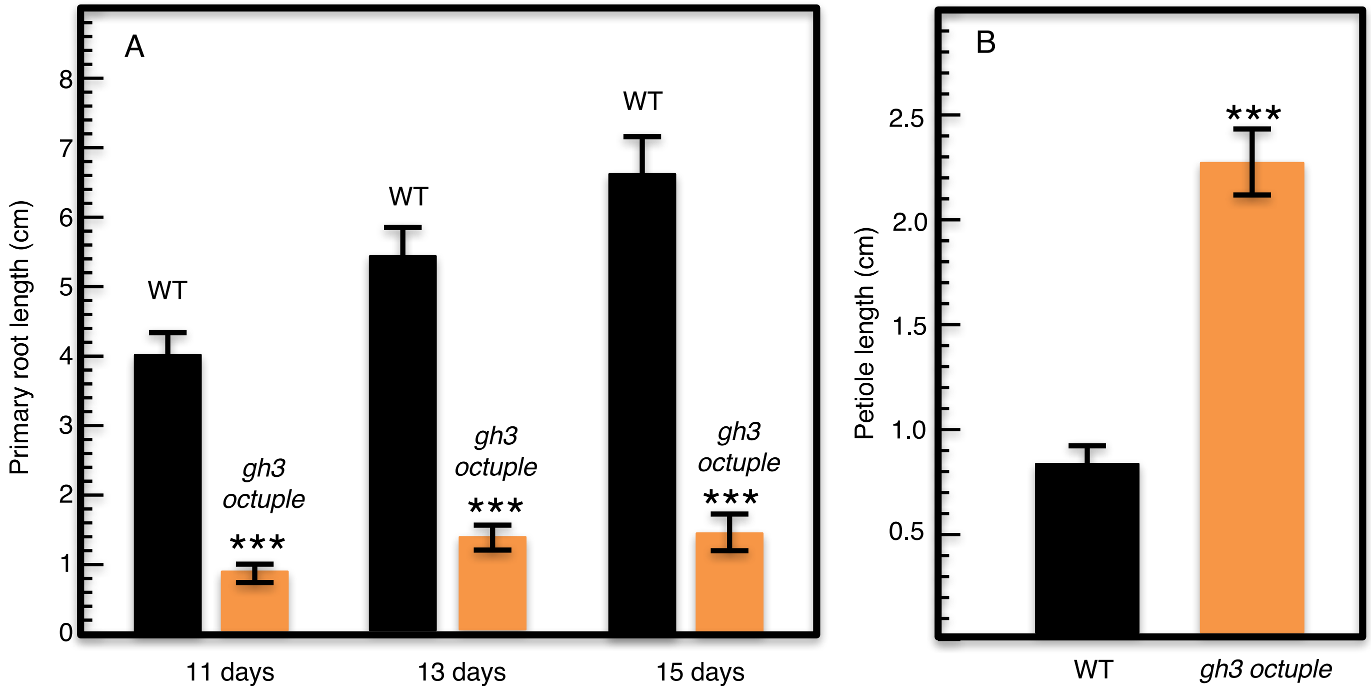


**SI Fig. 3. The impact of a lack of *GH3* genes on primary root and petiole development.** A) Both WT and *gh3 octuple* mutants grown on MS media on vertical plates. The *gh3 octuple* mutants had much shorter primary roots. B) Plants without group II *GH3* genes developed much longer petioles. Twenty seedlings of each genotype were measured for each trait. The results were subjected to student t-test. *** indicates < 0.001.

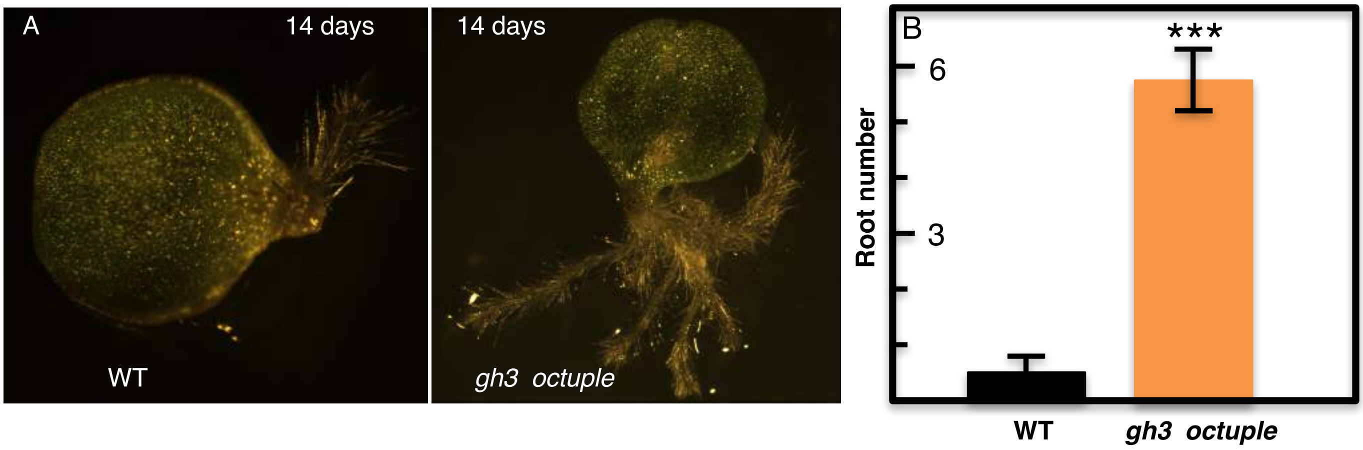


**SI Fig. 4. Explants of *gh3 octuple* mutants developed more and longer roots in hormone-free media**. A) Explants of WT and *gh3 octuple* mutants grown on MS media for two weeks. The mutant developed five roots whereas WT only had one root. B) On average, *gh3 octuple* explants developed at least 5 roots whereas WT under the same conditions only had fewer than one root.

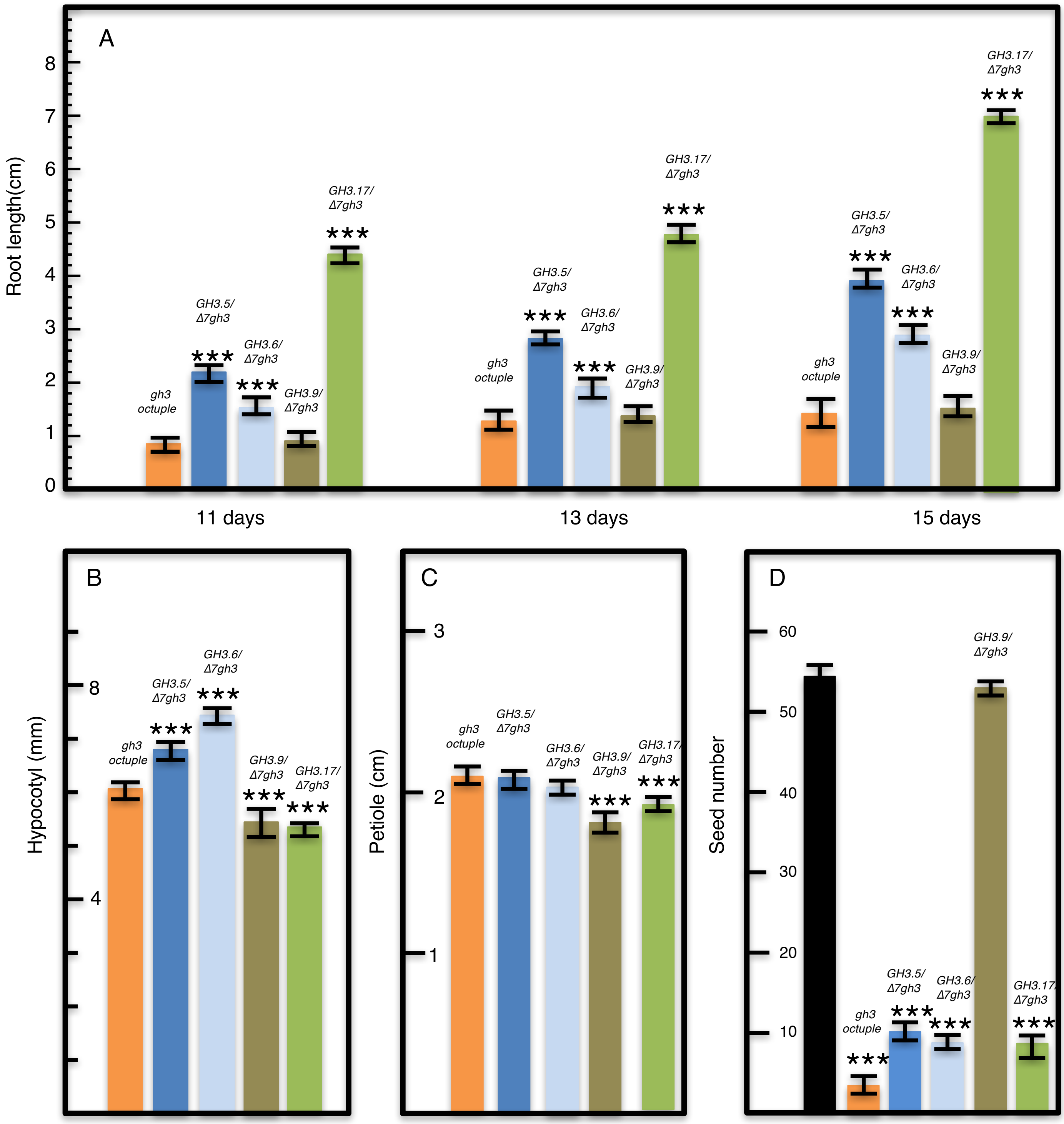


**SI Fig. 5. Suppression of *gh3 octuple* phenotypes by keeping one of the *GH3* genes.** A) The *GH3.5 Δ7gh3, GH3.6 Δ7gh3, GH3.17 Δ7gh3 septuple* mutants had longer roots than *gh3 octuple* mutants whereas *GH3.9 Δ7gh3* behaved similarly to *gh3 octuple.* B) Hypocotyl length of *gh3 octuple* and *gh3 septuple* mutants. Note that both *GH3.5 Δ7gh3* and *GH3.6 Δ7gh3* had longer hypocotyls than *gh3 octuple*. *C) GH3.9* and *GH3.17* appeared to have a role in petiole elongation. *D)* A single *GH3 (GH3.9)* was able to maintain Arabidopsis fertility. Seed number refers to seeds per silique. Twenty seedlings of each genotype were measured for each trait. For seed number measurements, twenty siliques of each genotype were analyzed. The results were subjected to student t-test. *** indicates < 0.001.

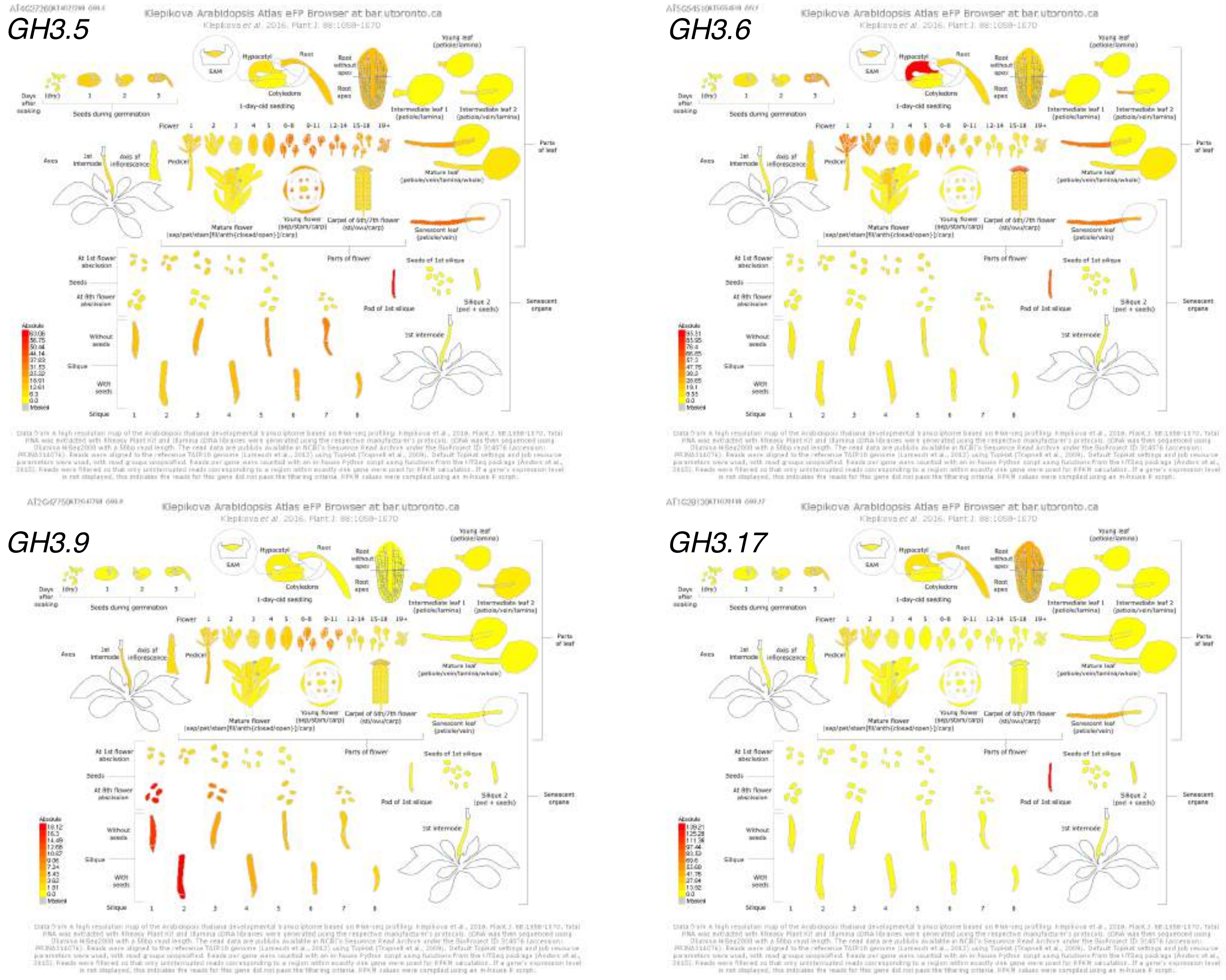


**SI Figure 6. The expression patterns of four *GH3* genes.** The results were from the Arabidopsis.org website. Note that *GH3.5*, *GH3.6* and *GH3.17* are expressed in roots whereas *GH3.9* has very low expressions in roots. Both *GH3.5* and *GH3.9* are expressed in siliques, but *GH3.9* appears to have higher expressions.

SI Table I. Genotyping primers used in this study.

| **Primer Name** | **Primer sequences (5' to 3')** |
|---|---|
| GH3.1-GT1 | TTTCTTCGCGTCAAGTTTTCCT |
| GH3.1-GT3 | ACTGGTCAAAACCGGTCGAG |
| GH3.1-GT2 | TCAAAGTAGACATGGTCCCTGG |
| GH3.2-GT1 | TGCTGACGTGGAACTTCTCC |
| GH3.2-GT3 | CGTCTCTTGAAGTGGTCGCT |
| GH3.2-GT2 | GGGCACATTGATTAACATTCCGT |
| GH3.3-GT1 | ACTTAGGCTCCGGGAACAAC |
| GH3.3-GT3 | GGGTTAAGTATATACCTTGTGAGGA |
| GH3.3-GT2 | GGTTGGTCTTAGGTGAAAGCA |
| GH3.4-GT1 | TGCAAAACTGAAAGTAACGAGACA |
| GH3.4-GT3 | ACAAGGGGATTGGTACTCCT |
| GH3.4-GT2 | GGCAGATCCATTTTTAAACCACAC |
| GH3.5-GT1 | ATCCGGTACAGGTTTTGACCA |
| GH3.5-GT3 | ACTCATCGCCCAAAAACAACT |
| GH3.5-GT2 | TTCTTTGTGGATGCGTCATTCT |
| GH3.6-GT1 | GCTCCCCGAATTCAAGATATGC |
| GH3.6-GT3 | AATGGTCAAACCTTGTGAGGA |
| GH3.6-GT2 | GGATGATTCTTGTTCAACTCTCGG |
| GH3.9-GT1 | GTGGCCCCCTTGTTGAGATT |
| GH3.9-GT3 | GCCTGACTTGCCCTTAGGAT |
| GH3.9-GT2 | CGCATCCAGGGAATCGAACC |
| GH3.17-GT1 | GAGTATAGTGAAGACCAAGTGAGT |
| GH3.17-GT3 | CCCCGAACTGCATCCAATA |
| GH3.17-GT2 | TGTCCCTGTTCCTGCATCTC |

Note: Mutants are genotyped using GT1/GT2 pair.  Zygosity of the mutants was determined using GT1/GT3 pair.
